# Characterization of the Gastrointestinal Tract Holstein x Angus Cross Cattle Microbiome During Harvest after Feed Withdrawal

**DOI:** 10.1101/2025.01.10.632417

**Authors:** M.K. Costello, J.C. McClure, J.A. Brown, H.C. Mantovani, S.C. Ricke

**Author notes:** Corresponding author: Dr. Steven C. Ricke, Department of Animal and Dairy Sciences, Office 2124 Meat Science and Animal Biologics Discovery Building, University of Wisconsin – Madison, 1933 Observatory Drive, Madison, WI 53706.

## Abstract

Stress during the beef pre-harvest period can induce an inflammatory response and acidotic conditions in the gastrointestinal tract (GIT), which affects the gastrointestinal tract microbiome. The objective of this study was to characterize the status of the GIT microbiome at harvest in beef cattle entering a small USDA processing facility. Nine beef cattle were shipped from a producer in Columbia County, WI to the USDA processing facility at University of Wisconsin-Madison and were harvested across four dates. Digesta samples were collected from eight GIT locations: rumen solids, rumen liquids, abomasum, duodenum, jejunum, ileum, cecum, and large intestines. After DNA extraction with the DNeasy Blood & Tissue Kit, the V4 region of the 16S rRNA gene was amplified and sequenced on the Illumina MiSeq platform. Sequences were analyzed for alpha and beta diversity metrics (ANOVA and ADONOS), core microbiome, ANCOM, and co-occurrence network analyses. Harvest date and GIT location had a significant impact on microbial diversity and community composition (P<0.05), and there was an interaction between GIT location and harvest date (P<0.05). Taxonomic composition shifted throughout the GIT, though *Prevotella* and *Treponema* were core members in several different GIT locations. The co-occurrence analysis revealed microorganisms potentially associated with clinical infections, such as *Moryella* in the rumen and *Acinetobacter* in the hindgut, were considered keystone species. These results suggest that the pre-harvest period may negatively impact the beef cattle GIT microbiome. Modulating the GIT microbiome during the pre-harvest period may offer an opportunity to improve food safety.

## Introduction

The pre-harvest process is a period in which cattle are transported from feedlots to processing facilities and held prior to slaughter. This is a stressful period for cattle, with several factors, such as transportation, temperature, stocking density, handling, and feed withdrawal, contributing to the stress response [1,2,3,4,5]. The compounding stress of these factors increases circulating hormones such as cortisol and glycoproteins such as haptoglobin and chromogranin A that deplete muscles glycogen stores, negatively affecting meat tenderness and color [2]. In addition to the effects on meat quality, the gastrointestinal tract (GIT) microbiome can be affected by several of these factors, including feed withdrawal [6,7]. Cattle typically undergo a varying feed withdrawal period during the pre-harvest process to reduce GIT contents during carcass dressing [8,9]. Despite the advantages during harvest, feed withdrawal has been associated with acidosis, increased fecal pathogen shedding, and inflammation [9]. Starvation during feed withdrawal diminishes populations of beneficial bacteria while encouraging the growth of acid-producing bacteria, such as *Streptococcus*, and pathogens, such as non-typhoidal *Salmonella* enterica or liver abscess-causing *Fusobacterium necrophorum* [6,7,10,11]. This poses major challenges for the beef industry as acidosis has been linked to liver abscesses and the shedding of fecal pathogens which can spread between animals [12,13]. Additionally, these conditions disrupt gut barrier function, which may contribute to the stress and inflammatory response [14,15]. As of late, there have been few studies measuring the impact of feed withdrawal on cattle during the pre-harvest process.

Of the studies examining the impact of the pre-harvest process and feed withdrawal on bovine health and food safety, few have focused on the entire GIT microbiome. As the whole GIT is exposed to the carcass during harvest, and each GIT compartment has a unique impact on host function, understanding the effects of these microbial communities offers an opportunity to partially mitigate food safety and meat quality concerns [9,16]. Despite considerable research into these effects, few studies have ventured beyond the rumen. Given the known relationship between the small intestine’s microbial communities and the stress and inflammatory responses, the small intestines are a prime target for modulating the stress from external factors [14,17]. In addition, the hindgut microbial communities influence food safety as fecal pathogen shedding is a major source of pathogen spreading and carcass contamination [18,19]. To begin parsing the impacts of each GIT microbial community during the pre-harvest process, this study aims to characterize the GIT microbiome of beef cattle at harvest and begin identifying potential opportunities to improve carcass quality and food safety.

## 1. Materials & Methods

### Sample Collection

Angus-Holstein cross cattle were shipped from a single producer based in Columbia County, WI around 20 months of age to the University of Wisconsin-Madison (UW-Madison) USDA processing facility in the Meat Science & Animal Biologics Discovery building. Animals were fed a grassy hay and corn diet mixed with the Gain Master 55:35 RT #1707 pellet (Big Gain Inc., Mankato, MN) at a 5% inclusion rate. Cattle were withheld feed overnight, for approximately 10 hours prior to harvest, and were transported to UW-Madison the morning of harvest. Nine animals were harvested on 4/6/23 (n=2), 4/25/23 (n=2), 6/8/23 (n=3), and 8/10/23 (n=2) according to USDA specifications. Harvest began at approximately 7:30 AM each morning, and the average temperatures the week before harvest were as follows: 8.89 C on 4/6/23, 11.67 C on 4/25/23, 22.22 C on 6/10/23, and 22.78 C on 8/10/23. Digesta content samples were taken post-evisceration from seven locations throughout the GIT: rumen, abomasum, duodenum, jejunum, ileum, cecum, and large intestines. Due to the low volume of digesta content throughout the GIT, samples were collected based on visual identification of the compartments, shown in **Figure 1**. Immediately following collection, rumen samples were separated into solid and liquid fractions using four layers of bleached cheesecloth (Grainger Industrial Supply Lake Forest, IL, USA). Samples were stored in 50 mL conical tubes (Eppendorf, Hamburg, Germany) at -20° Celsius before DNA extractions.

**Figure 1.**
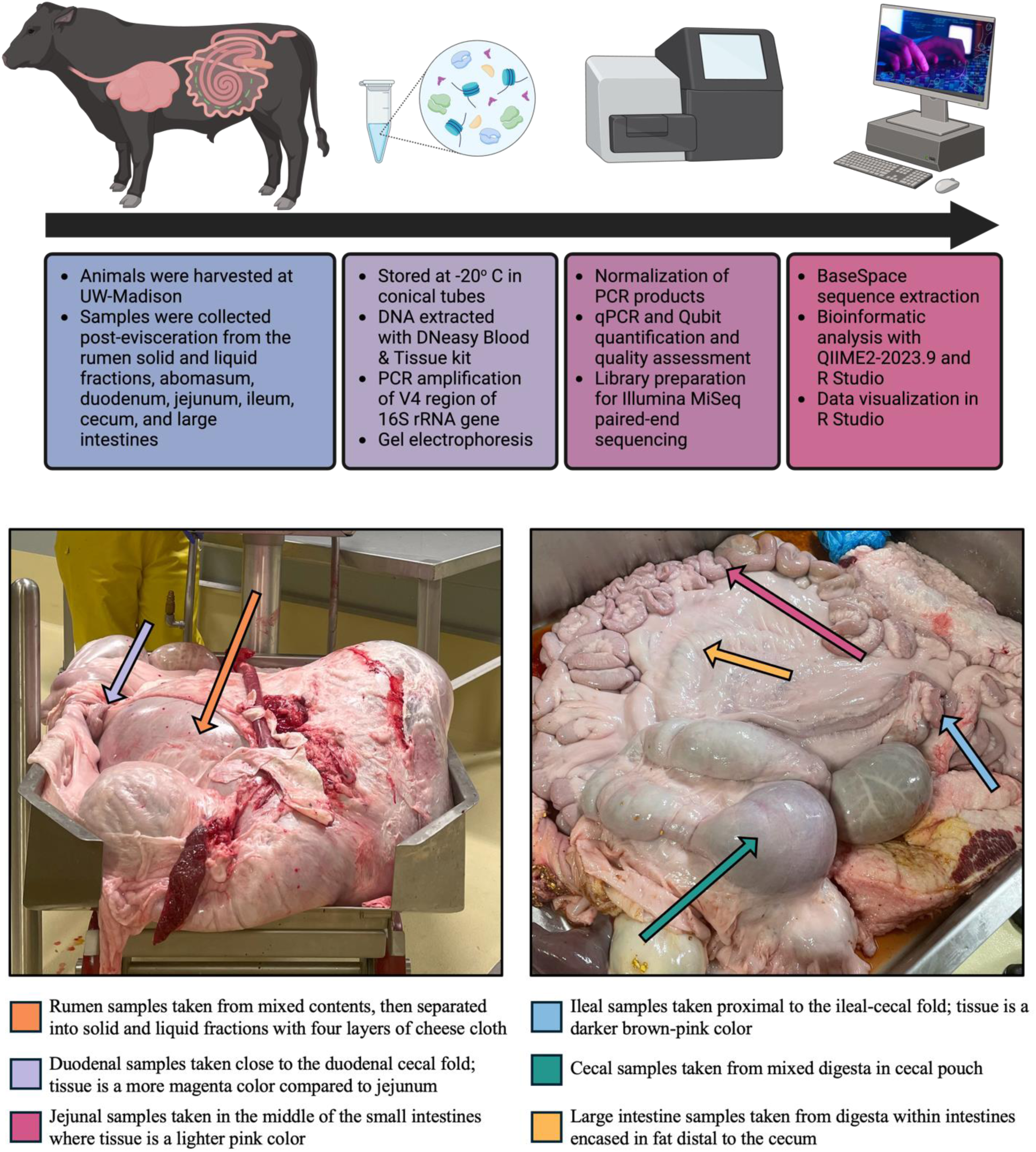
Experimental overview and sampling techniques. Figure made with Biorender (Agreement number: ML27HFY3SN).

### 16S rRNA Gene Sequencing

Before DNA extractions, frozen samples were thawed, and 325 mg were aliquoted into 2 mL microcentrifuge tubes (Eppendorf, Hamburg, Germany). A Qiagen DNeasy Blood & Tissue Kit (Qiagen, Hilden, Germany) was used to extract DNA from the samples with a 10-minute bead beating step. DNA concentrations were quantified with a Nanodrop^TM^ 1000 spectrophotometer (Thermo Fisher Scientific, Waltham, MA, USA), and samples with DNA concentrations above 15 ng/μL were diluted to 10 ng/μL in AE buffer, while samples with concentrations below 15 ng/μL were not. Following DNA extraction and dilution, the V4 region of the 16S rRNA gene region was amplified with a high-fidelity polymerase (Accuprime Pfx DNA polymerase, Thermo Fisher Scientific, Waltham, MA, USA) and dual-indexed primers, with eight nucleotide barcode sequences as developed by [20]. The PCR products were confirmed using gel electrophoresis, and successfully amplified products were normalized to 20 nM with a SequalPrep^TM^ Normalization kit (Life Technologies, Carlsbad, CA, USA). Final libraries were created with 5 μL of each of the normalized samples. The final library DNA concentrations were determined with an Illumina platform-specific KAPA library quantification kit (Kapa Biosystems, Inc., Wilmington, MA, USA) and 1x High Specificity Assay kit on a Qubit 4 fluorometer (Invitrogen, Carlsbad, CA, USA). The pooled library was then diluted to 20 nM and combined with HT1 buffer, 20 nM of PhiX v3 control, and 0.2 N NaOH to provide a final concentration of 6 pM. The sample solution was mixed with PhiX control v3 (20% v/v) before 600 μL were loaded into a MiSeq v2 (500 cycles) reagent cartridge (Illumina, San Diego, CA, United States).

The sequences were downloaded from Illumina BaseSpace. The sequences were subsequently downloaded locally and input into QIIME2-amplicon-2023.9 via the Casava1.8 paired-end pipeline [21]. Amplicon sequencing variant (ASV) taxonomic assignment was completed with classify-sklearn and the SILVA 2023.9 database with a confidence limit of 95%. After visualization, the sequences were trimmed with DADA2 in the chimera consensus pipeline [22]. The taxonomic output file, sample metadata, rooted phylogeny tree, and feature table were uploaded into R Studio for further statistical analyses.

### Statistical Analyses

Several packages were used for statistical analysis and visualization, including Phyloseq [23], qiime2R [24], DEseq2 [25], vegan [26], microbiomeutilities [27], and SpiecEasi [28]. Using linear regression models, alpha diversity was analyzed and visualized for diversity and richness with Pielou’s Evenness, Observed Features, Simpson’s Index, Chao Index, and Shannon’s Diversity Index. These models were assessed for normality with the Shapiro Test, the Lillie Test, Cramér–von Mises test, the Anderson-Darling test, and Akaike’s information criterion (AICcmodavg) to find the model of best fit before an analysis of variance (ANOVA) test was completed to determine group significance and interactions. Alpha diversity pairwise comparisons were analyzed using Tukey’s Honest Significant Differences test. Beta diversity was assessed with two quantitative indicators, the Bray-Curtis dissimilarity index and the Weighted Unifrac distance matrix, considering both variation and population dispersion with the Analysis of Similarity (ANOSIM) function. Core microbial members were identified with a core microbiome analysis (microbiomeutilities) with a detection setting of 0.01 and a prevalence of 20% due to high sample variability [27]. Differential abundance analyses were completed with the DESeq2 package, which uses an analysis of the compositional profiles of microorganisms (ANCOM) and a Wald test at 0.01 [25]. Co-occurrence networks were determined and visualized with methods described by Amorín de Hegedüs et al. [29] and the mdmnets package [30]. Due to the low sample size, GIT locations within a compartment were analyzed together for the differential abundance analysis and the co-occurrence network analysis. The rumen (rumen solids and rumen liquids), the small intestines (duodenum, jejunum, and ileum), and the hindgut (cecum and large intestines) were pooled to create 3 locations for these analyses. Data were visualized with ggplot2 and RColorConesa.

### Data Availability

The raw data were deposited in the NCBI BioProject database (Accession Number: PRJNA1204970)

## 2. Results

### Microbial diversity varied across the GIT and between harvest date

In this study, we aimed to characterize the microbiome throughout the GIT at harvest. The effects of GIT location, harvest date, and the interaction between both variables on sample evenness and richness were assessed with ANOVA (Table 1), and the phylogenetic diversity and abundances of each sample were determined with ANOSIM considering these factors and their combinatory effect (Table 1). Gastrointestinal tract location was the strongest indicator of microbial diversity (P<0.05; Table 1; Figures 2A-D and community dissimilarity (P<0.05 Table 1; Figures 3A and 3B). The cecum had the highest richness and evenness compared to the other GIT locations, while the jejunum had the lowest richness (P<0.05; Figures 2A-D). Harvest date also had a significant impact on sample diversity and richness (P<0.05; Table 1; Figures 2E-H) and community composition (P<0.05; Table 1; Figures 3C-D). Cattle harvested on 4/25/23 (n=2) had the highest microbial diversity and significantly higher richness and evenness than cattle harvested on 8/10/23 (n=2) (P<0.05; Figures 2E-H). Community composition, sample richness, and evenness were also impacted by an interaction between GIT location and harvest date (Table 1). Each GIT compartment exhibited a distinct community structure (P < 0.05), though the GIT locations within each compartment did not (P<0.05). Therefore, the sampling locations within each GIT compartment were pooled for some analyses.

**Figure 2.**
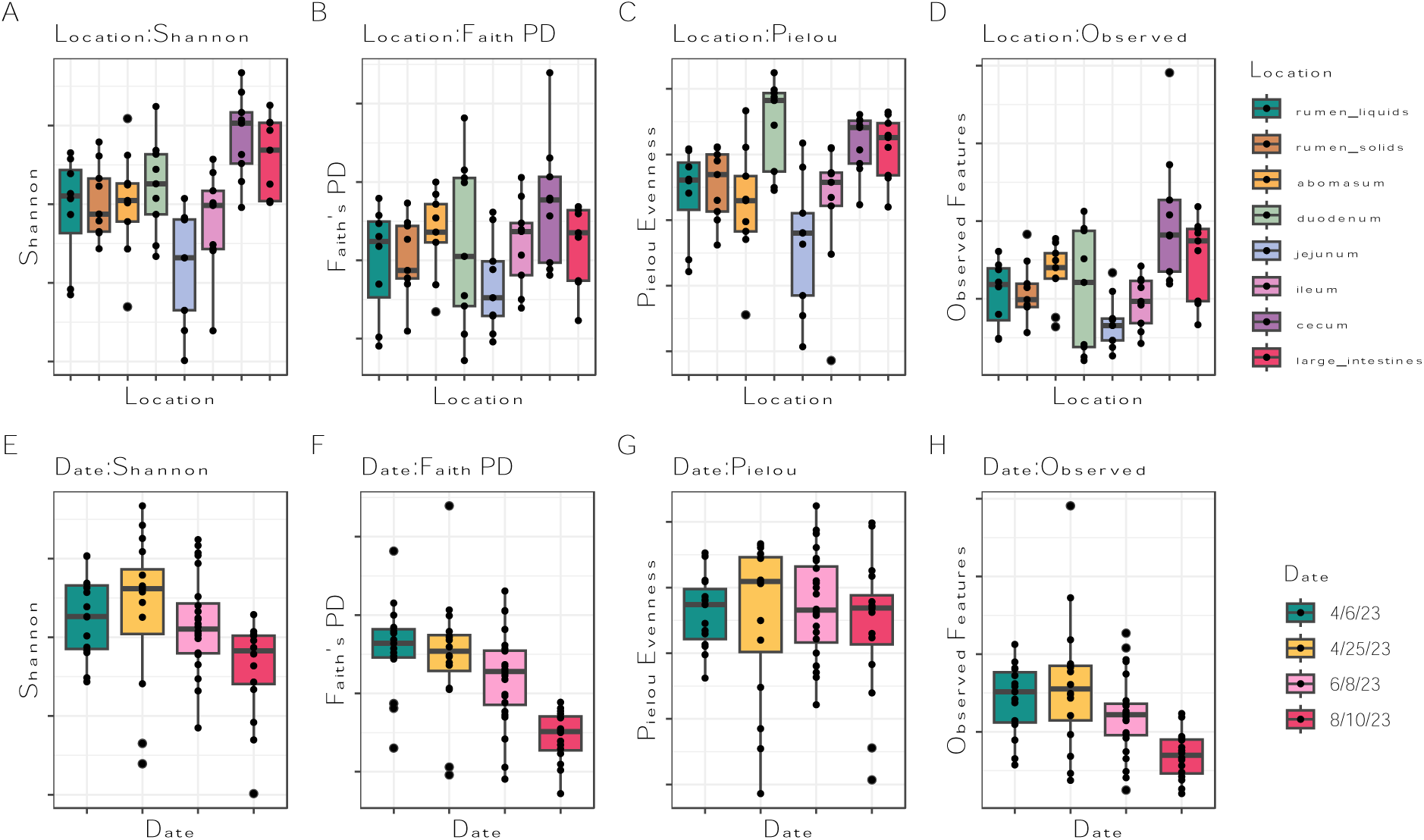
Microbial diversity (A, E: Shannon’s Diversity; B, F: Faith’s Phylogenetic Diversity; C, G: Pielou’s Evenness; D, H: Observed Features) between location (A, B, C, D) and harvest date (E, F, G, H).

**Figure 3.**
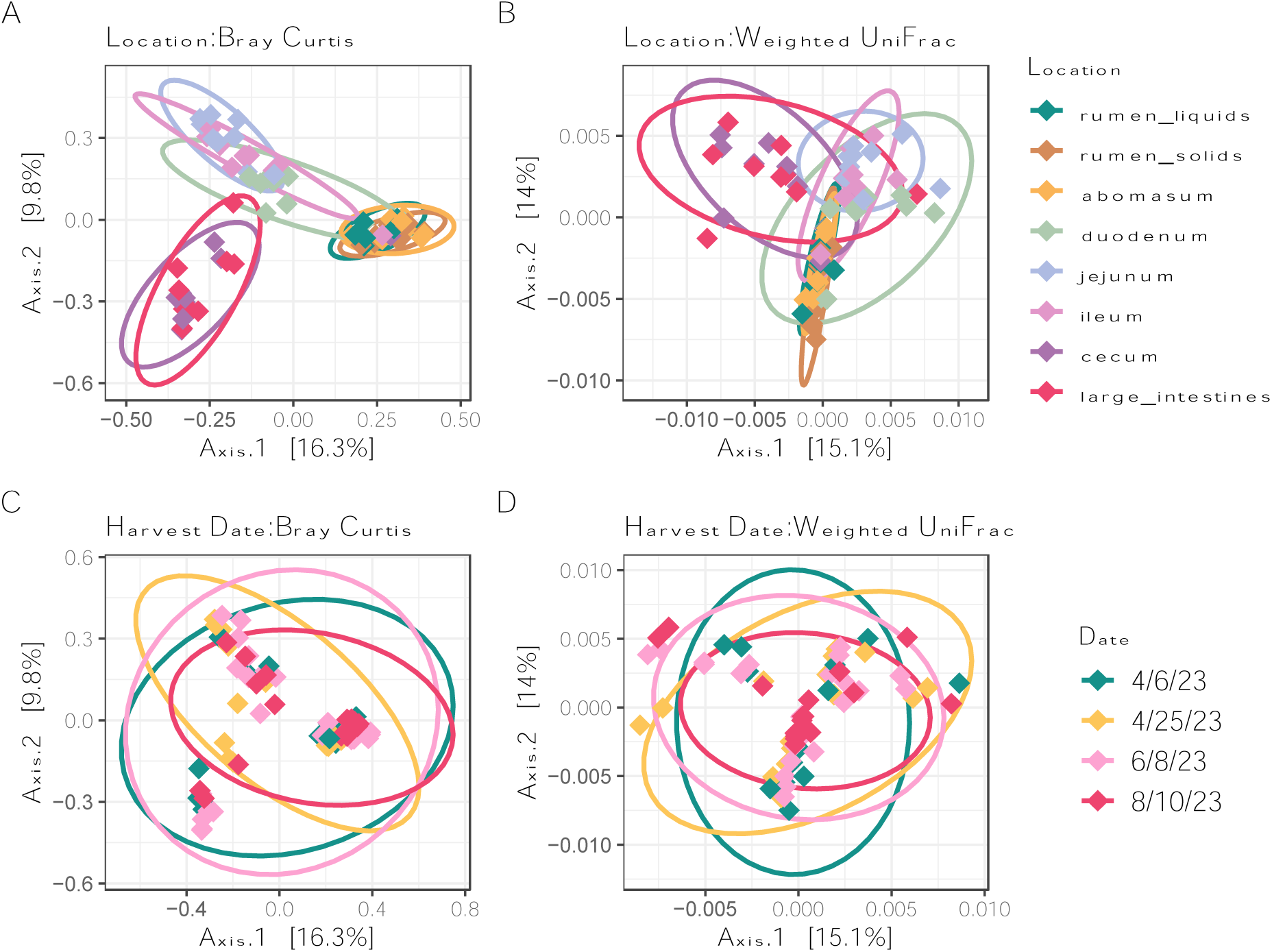
Community dissimilarity (A, C: Bray-Curtis; B, D: Weighted Unifrac) between GIT location (A, B) and harvest date (C, D).

**Table 1.** ANOVA results for each alpha diversity metric and ADONIS results for each beta diversity metric.

| Alpha Diversity |  |  |  |  |  |  |
| --- | --- | --- | --- | --- | --- | --- |
| Factor | Shannon | Simpson | Chao1 | Observed | Pielou | Faith |
| Date | 0.0004* | 0.11 | 0.000000239* | 0.000000241* | 0.51 | 0.000000309* |
| Location | 0.00000148* | 0.0026** | 0.000000218* | 0.000000234* | 0.0059** | 0.00000169* |
| Date*Location | 0.053 | 0.32 | 0.0454* | 0.04* | 0.059 | 0.0025* |
| Beta Diversity |  |  |  |  |  |  |
| Factor | Bray | Jaccard | Unweighted Unifrac | Weighted Unifrac |  |  |
| Date | 0.01* | 0.01* | 0.001* | 0.001* |  |  |
| Location | 0.01* | 0.01** | 0.001** | 0.001* |  |  |
| Date*Location | 0.04 | 0.07 | 0.15 | 0.114 |  |  |

\* P<0.05

### Community composition differed across the GIT

Community structure shifted across the GIT, with many shared taxa between neighboring GIT locations (Figures 4 and 5). A top 40 most abundant taxa analysis was completed alongside a core microbiome analysis, with a prevalence of 50% and 0.1% detection, to determine microbial community structure (Supplementary Tables 1 and 2). The rumen liquid and solid fractions had similar top taxa and core members. *Prevotellaceae* YAB2003*, Prevotella, Succinivibrionaceae* UCG-001*, Treponema, Muribaculaceae,* and *Rikenellaceae* RC9 gut group were among the most abundant in the rumen liquids and solids. These listed taxa were also shared core members between the rumen liquids and solids (Figure 5). There was minimal methanogen representation in the rumen, with only *Methanobrevibacter* as the 23^rd^ most abundant taxa in the rumen solids fraction. The top taxa and core members in the rumen were nearly identical in the abomasum and included *Prevotella, Succinivibrionaceae* UCG-001, and *Treponema*.

**Figure 4.**
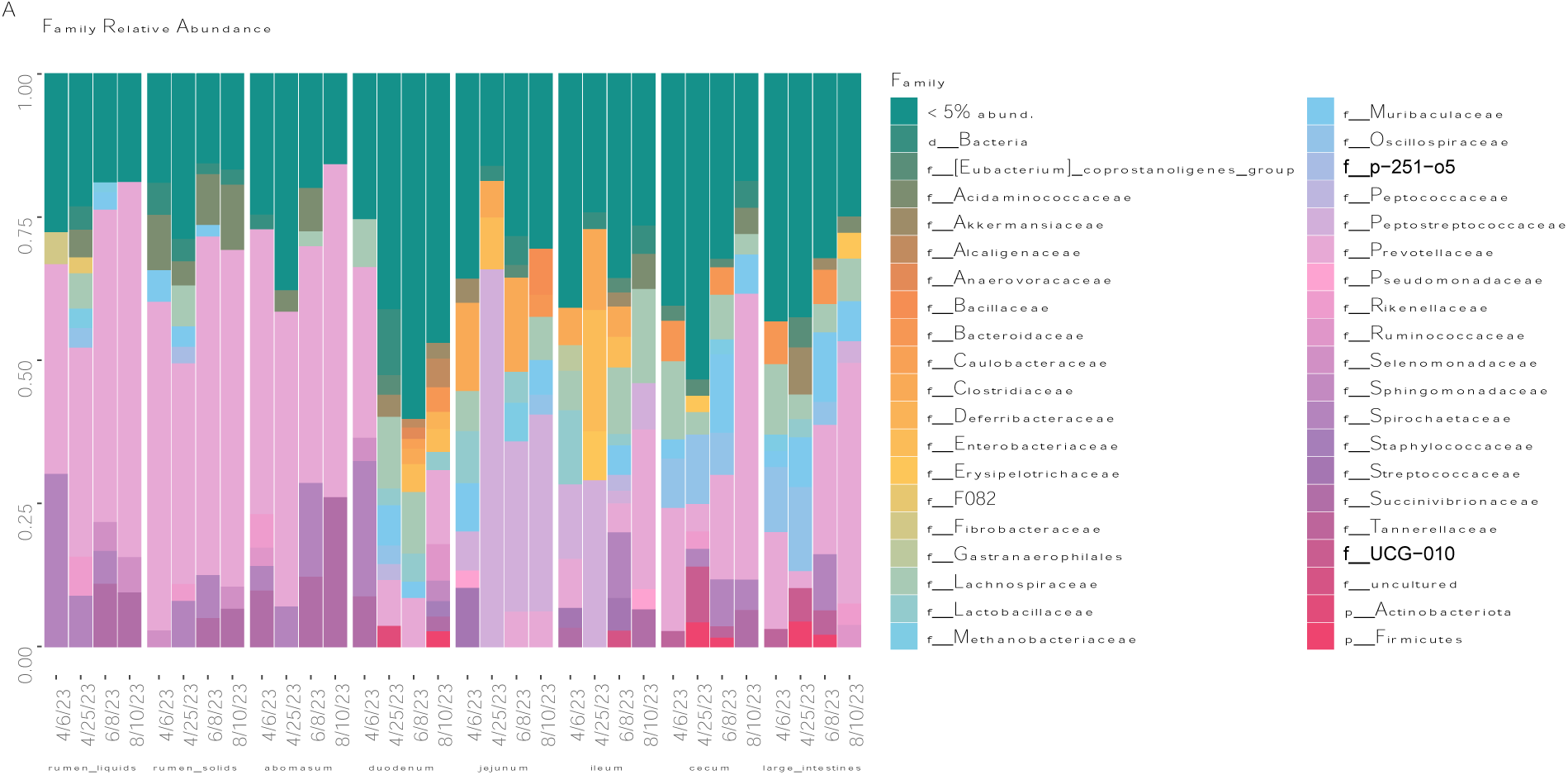
Taxonomic bar plots of family level relative abundances.

**Figure 5.**
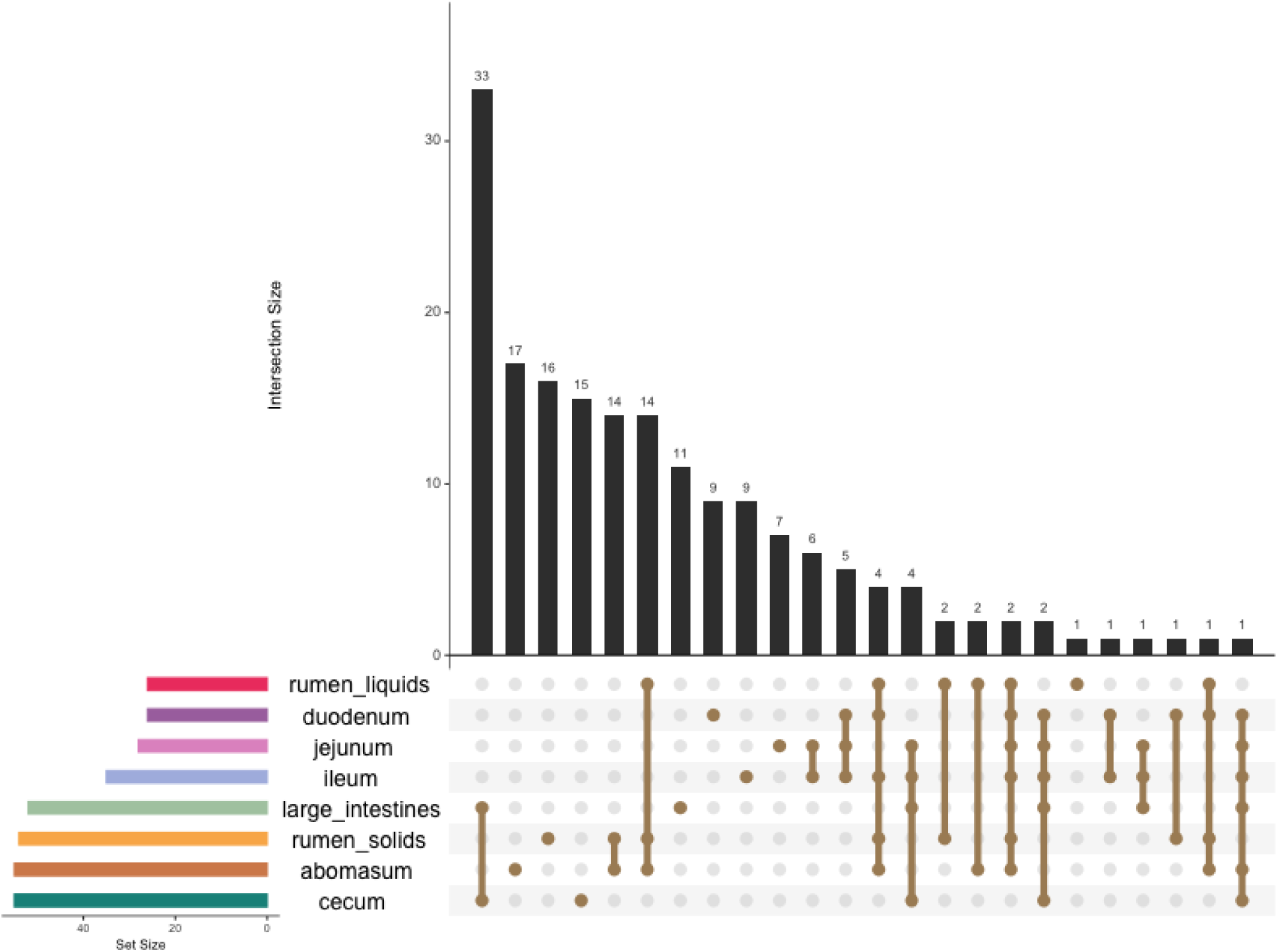
Shared ASVs between GIT compartments. Amplicon sequencing variants were considered core members if they were higher than 0.01% abundance and detected in at least 50% of the samples. The total number of core members in each location are shown on the left and the number of shared ASVs are visualized as intersection size.

The lower GIT shared numerous highly abundant and core taxa. In the duodenum, *Prevotella, Lachnospiraceae, Muribaculaceae, Pseudomonas,* and *Lactobacillus* were considered core members (Supplementary Table 2; Figure 5). Several bacteria, *Akkermansia. Peptostreptococcaceae*, *Clostridium senso stricto 1, Lactobacillus,* and Enterobacteriaceae, were identified as the most prevalent taxa in the jejunum and ileum. While not a core member, *Staphylococcus* was represented in the top forty most abundant taxa in the jejunum and ileum (Supplementary Table 1). The duodenum, jejunum, and ileum shared many core taxa, including *Turicibacter*, *Methanobrevibacter, Akkermansia,* and *Clostridium senso stricto* 1. As observed in the small intestines, the cecum and large intestines shared many top taxa, including *Alloprevotella, Prevotella,* and *Clostridia.* There were 55 core members in the cecum, including *Methanobrevibacter, Clostridium senso stricto* 1, *Muribaculaceae*, and *Oscillospiraceae* UCG-005 (Supplementary Table 2; Figure 5). The large intestines shared many of its 52 core taxa with the cecum, except for *Treponema, Succinivibrio,* and several *Alloprevotella* taxa. In contrast to the rumen fractions, two methanogens were among the top forty most abundant taxa in the hindgut, including *Methanobrevibacter* and *Methanocorpusculum*.

### Low abundant taxa were critical to community structure in each GIT compartment

Co-occurrence networks were built for each GIT compartment: the rumen (rumen solids and rumen liquids; Figure 6A), small intestines (duodenum, jejunum, and ileum; Figure 6B), and hindgut (cecum and large intestines; Figure 6C). A co-occurrence network is analyzed and assessed with the determination of three factors: degree of centrality, closeness centrality, and betweenness centrality [29]. Hub scores are assigned, taking into consideration all three of these metrics, and a hub score of 1 indicates a keystone member in an ecosystem. All 25 of the highest hub scores in the rumen and small intestines had hub scores above 0.7 (Table 2). However, 19 out of the 25 highest-scoring hubs were below 0.7 in the hindgut. *Moryella* was considered a keystone species (1.00) in the rumen despite having relatively low abundance in both fractions, and *Megasphaera* was the second highest scoring hub (0.963). In the small intestines, *Muribaculaceae* had the highest scoring hub score (1.00). While several *Muribaculaceae* taxa were considered core members across the locations within the small intestines, the taxa identified as keystone members were not. Three core members in the small intestines, *Methanobrevibacter* (0.778), *Micrococcaceae* (0.928), and *Prevotella* (0.962), were represented in the top 25 highest-scoring hubs in the small intestines, alongside several less predominant taxa, including *Eubacterium coprostanoligenes* group (0.831) and *Streptococcus* (0.829). Interestingly, *Acinetobacter* (1.00) was the keystone species in the hindgut despite not being in the top 40 most abundant taxa of the cecum or large intestines, with no representation in the core microbiome analysis. *Christensenellaceae* R7 group (0.782)*, Pyramidobacter* (0.729), *Roseburia* (0.649), and *Defluvitaleaceae* UCG-001 (0.645) were among the least abundant genera in the top 25 highest hub score groups in the hindgut.

**Figure 6.**
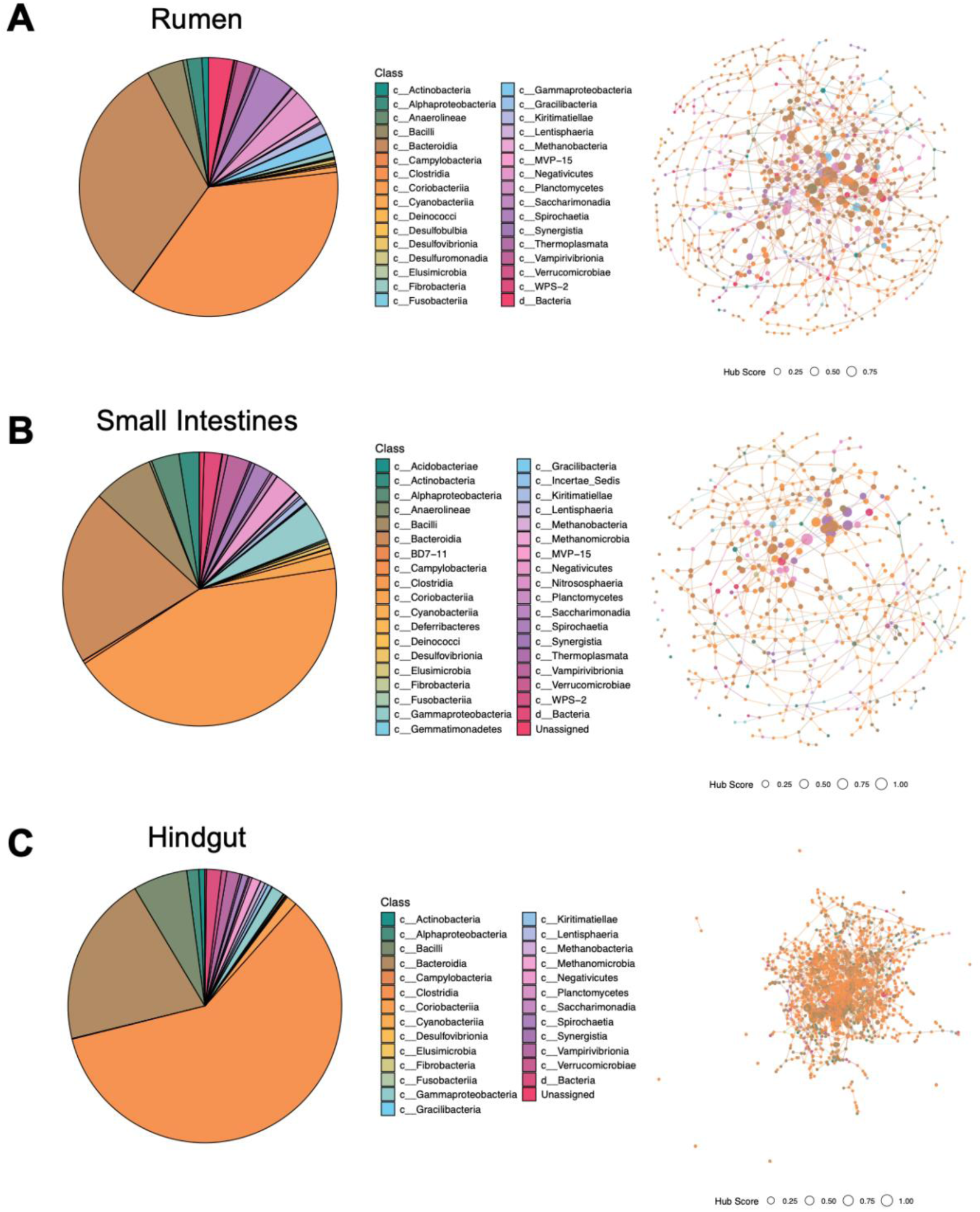
Co-occurrence networks for the rumen (rumen solids and liquids) (A), small intestines (duodenum, jejunum, and ileum) (B), and hindgut (cecum and large intestines) (C). The distribution of class in the co-occurrence networks are visualized in the pie charts. Hub networks at the class level are shown for each compartment. High scoring hubs are represented by a larger dot while low scoring hubs are represented by a smaller dot, as shown in the legend.

**Table 2.** Top 25 highest scoring hubs in each GIT compartment: foregut (rumen solids and liquids), small intestines (duodenum, jejunum, and ileum), and hindgut (cecum and large intestines).

| Compartment | Hub Score | Genus |
| --- | --- | --- |
| rumen | 1.000 | *Moryella* |
| rumen | 0.963 | *Megasphaera* |
| rumen | 0.919 | *Clostridia* |
| rumen | 0.890 | *Lachnospiraceae* |
| rumen | 0.878 | *Clostridia* |
| rumen | 0.851 | *Lachnospiraceae_NK3A20_group* |
| rumen | 0.846 | *Prevotella* |
| rumen | 0.831 | *Prevotellaceae_YAB2003_group* |
| rumen | 0.829 | *Prevotella* |
| rumen | 0.815 | *Lachnospiraceae* |
| rumen | 0.813 | *Lachnospiraceae_NK3A20_group* |
| rumen | 0.793 | *Lachnospiraceae* |
| rumen | 0.790 | *Prevotella* |
| rumen | 0.781 | *Bacteroidales* |
| rumen | 0.779 | *Proteobacteria* |
| rumen | 0.778 | *Rikenellaceae_RC9_gut_group* |
| rumen | 0.771 | *Prevotellaceae* |
| rumen | 0.771 | *Bacteroidales* |
| rumen | 0.760 | *Spirochaetaceae* |
| rumen | 0.758 | *Bacteroidales* |
| rumen | 0.754 | *Elusimicrobium* |
| rumen | 0.750 | *Oribacterium* |
| rumen | 0.729 | *Muribaculaceae* |
| rumen | 0.726 | *Prevotellaceae_UCG-004* |
| rumen | 0.726 | *Lachnospiraceae* |
| small intestines | 1.000 | *Muribaculaceae* |
| small intestines | 0.962 | *Prevotella* |
| small intestines | 0.933 | *Lachnospiraceae* |
| small intestines | 0.928 | *Micrococcaceae* |
| small intestines | 0.879 | *NK4A214_group* |
| small intestines | 0.864 | *Bacteroidales* |
| small intestines | 0.831 | *[Eubacterium]_coprostanoligenes_group* |
| small intestines | 0.829 | *Streptococcus* |
| small intestines | 0.823 | *Prevotella* |
| small intestines | 0.821 | *Lachnospiraceae* |
| small intestines | 0.814 | *Akkermansia* |
| small intestines | 0.805 | *Bacteroides* |
| small intestines | 0.798 | *Prevotella* |
| small intestines | 0.796 | *F082* |
| small intestines | 0.787 | *Prevotella* |
| small intestines | 0.784 | *Enterobacterales* |
| small intestines | 0.784 | *Chloroplast* |
| small intestines | 0.781 | *UCG-005* |
| small intestines | 0.778 | *vadinBE97* |
| small intestines | 0.778 | *Methanobrevibacter* |
| small intestines | 0.777 | *Lachnospiraceae* |
| small intestines | 0.776 | *F082* |
| small intestines | 0.768 | *Lachnospiraceae* |
| small intestines | 0.767 | *Mitochondria* |
| small intestines | 0.760 | *Bacteroides* |
| Hindgut | 1.000 | *Acinetobacter* |
| Hindgut | 0.864 | *Lachnospiraceae* |
| Hindgut | 0.855 | *UCG-005* |
| Hindgut | 0.782 | *Christensenellaceae_R-7_group* |
| Hindgut | 0.729 | *Pyramidobacter* |
| Hindgut | 0.724 | *Lachnospiraceae* |
| Hindgut | 0.692 | *Clostridia_UCG-014* |
| Hindgut | 0.678 | *Muribaculaceae* |
| Hindgut | 0.674 | *Clostridia* |
| Hindgut | 0.664 | *Enterobacterales* |
| Hindgut | 0.656 | *Peptostreptococcaceae* |
| Hindgut | 0.654 | *UCG-010* |
| Hindgut | 0.648 | *Gastranaerophilales* |
| Hindgut | 0.646 | *Roseburia* |
| Hindgut | 0.645 | *Defluviitaleaceae_UCG-011* |
| Hindgut | 0.640 | *Lachnospiraceae* |
| Hindgut | 0.640 | *Lactobacillus* |
| Hindgut | 0.637 | *Clostridium_sensu_stricto_6* |
| Hindgut | 0.637 | *UCG-010* |
| Hindgut | 0.632 | *Lactobacillus* |
| Hindgut | 0.629 | *[Eubacterium]_coprostanoligenes_group* |
| Hindgut | 0.623 | *UCG-009* |
| Hindgut | 0.621 | *Lachnospiraceae* |
| Hindgut | 0.592 | *Peptostreptococcaceae* |
| Hindgut | 0.589 | *Prevotella* |

### Harvest date significantly impacted the GIT microbial communities in each compartment

Harvest date affected community structure within GIT locations. The differences between the most diverse harvest date, 4/25/23, and the least diverse harvest date, 8/10/23, were analyzed by GIT compartment (Figure 7A-D). These results are shown in Figure 7A-D. In the foregut, several *Prevotella* ASVs, *Methanobrevibacter, Fibrobacter*, and *Bacteroidota* were enriched on 8/10/23. There were 39 differentially abundant taxa in the small intestines, including many classified as family *Prevotellaceae, Lachnospiraceae,* and *Rikenellaceae.* There were three methanogenic ASVs enriched in the small intestines on 4/25/23, two classified as *Methanobrevibacter* and *Methanocorpusculum.* These taxa were also enriched in the hindgut on 4/25/23 along with *Eubacterium coprostanoligenes* group, *Clostridia* UCG-014, and *Desulfovibrio*.

**Figure 7.**
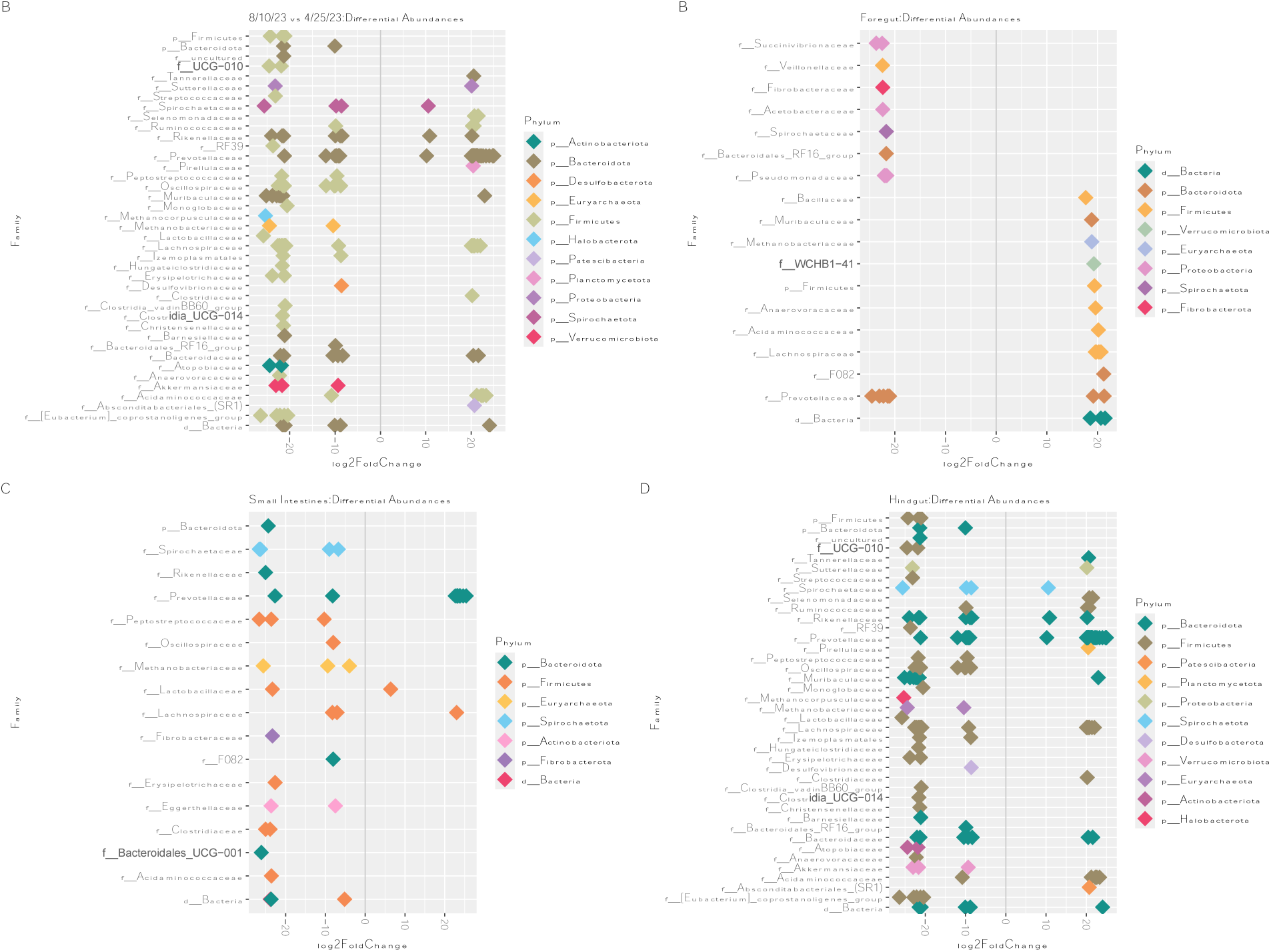
Differential abundance analysis of 8/10/23, the least diverse sampling date, vs. 4/25/23, the most diverse sampling date, on a log-fold change scale. A negative log fold change indicates taxa enriched on the 4/25/23 timepoint while a positive log fold change is enriched on 8/10/23. These two harvest dates had the most significant differences in microbial diversity (P=0.0058). Taxa were considered differentially abundant at P<0.01. An analysis was performed on all compartments (A), the rumen solids and liquids (B), the small intestines (duodenum, jejunum, and ileum) (C), and the hindgut (cecum and large intestines) (D).

## 3. Discussion

The GIT microbiome is central to beef cattle production and food safety [8,31]. The rumen and fecal microbiome have been tied to the prevention of pathogen colonization and host immune response [31]. Under normal conditions, the collective rumen microbiome prevents the rapid growth and domination of harmful bacteria, such as *Salmonella* or *F. necrophorum* [6,7]. However, during starvation conditions, beneficial bacteria die off and opportunistic pathogens rapidly proliferate [12]. The small intestines and hindgut microbial communities function similarly without stressors, harboring populations of bacteria, such as *Succinivibrio, Prevotella,* and *Butyrivibrio,* that promote mucus secretion, blocking pathogenic colonization [32,31]. The absence of these beneficial bacteria can shift dominance to mucus-degrading communities that stimulate inflammatory gene expression in the gut epithelium, contributing to systemic inflammation [17,33]. In the hindgut, acidosis can damage the epithelium and promote pathogen proliferation [13]. Furthermore, bacteria that are normally beneficial may become functionally harmful during dysbiosis. For instance, certain *Clostridium* and *Escherichia* species may occupy a small niche during high substrate availability, becoming opportunistic when competing bacteria are eliminated [34,35]. A healthy and responsive GIT microbiome is important during pre-harvest transportation and lairage period to reduce adverse effects from stress and feed withdrawal [36,31].

The current study surveyed the GIT microbiome of nine beef cattle entering a small USDA processing facility from a single producer on four different dates. The GIT microbiome varied between harvest dates, as shown by the significant differences in microbial diversity and community composition. For instance, cattle harvested during the first three harvest dates had significantly higher microbial diversity than cattle harvested on 8/10/23, and there were upwards of 100 differentially abundant taxa between the most diverse harvest date, 4/25/23, and the least diverse harvest date in the hindgut. The low number of animals harvested on each date prevents broad conclusions, however, there are numerous factors that may have contributed to the observed differences between harvest dates. An overnight feed withdrawal period, which was estimated to be around 10 hours was reported by the producer. Given the USDA processing facility’s overnight feed withdrawal guideline and the lack of strict requirements, it is possible that some of the feed withdrawal estimations varied by several hours. In addition to inconsistent feed withdrawal periods, there were major differences in temperature between harvest dates. The first two harvest dates, 4/6/23 and 4/25/23 had lower average weekly temperatures than the latter two harvest dates, 6/8/23 and 8/10/23. It has been previously reported that both feed withdrawal and high temperatures negatively affect the GIT microbiome [6,7,34]. Both conditions increase the abundance of lactic acid producing bacteria in the rumen, which lowers ruminal pH and disrupts regular gut barrier function [6,7,34]. Thus, more studies are necessary to explore the effects of different pre-harvest factors on the GIT microbiome and compare the microbial effects to food safety and animal welfare concerns.

The pre-harvest GIT microbiome networks and composition harbored taxa related to inflammation. *Megasphaera*, a lactic acid-utilizing bacteria, and *Streptococcus*, a lactic-acid-producing bacteria, were high scoring hubs in the rumen, which may contribute to acidosis [37,6]. Ruminal acidosis can directly affect the inflammatory response as the osmotic pressure from acidic contents can damage the rumen epithelium, promoting inflammatory gene expression [38,39]. Additionally, the antagonistic association between acidic conditions and bacterial diversity and richness can promote pathogen proliferation [40,41]. The enriched abundance of certain pathogens can lead to an accumulation of bioamines and lipopolysaccharides that can further aggravate the epithelium [40,41]. In the small intestines, *Streptococcus* and *Muribaculaceae* were high scoring hubs as well as core members. *Muribaculaceae* is a known mucus degrader and increases in abundance during fasting periods [42,43]. This trend continued through the large intestines, where *Muribaculaceae* and *Parabacteroides* were core members. While typically reported as a commensal taxon, *Parabacteroides* has been associated with disease and chronic inflammation in humans under stress conditions [32,44]. Meat quality is negatively affected by increased inflammation as inflammation can elevate cortisol levels [45,2]. As animals are simultaneously experiencing increased stress from external pre-harvest factors, the compounded inflammation from the GIT only exacerbates meat quality concerns. Therefore, pro-inflammatory taxa that affect gut barrier function may increase pathogen migration from the GIT [46].

Several potentially pathogenic and spoilage bacteria were identified throughout the GIT. *Moryella,* the highest hub score in the rumen and a keystone genus in the community network has previously been associated with mastitis and *E. coli* O157:H7 fecal shedding in dairy cattle [47,3]. The association between *Moryella* and pathogenesis is likely due to its involvement in indole production, a signaling molecule that may affect bacterial virulence and LPS production, which contributes to host inflammation [48,47]. *Moryella* can also migrate from the GIT into pus, which may exacerbate the stress response by penetrating the gut barrier in compromised pre-harvest animals [47]. The second most abundant taxa in the small intestines, *Clostridium senso stricto 1* (a taxonomic group including *Clostridium perfringens*), has been associated with diarrhea in calves [4]. Additionally, *Pseudomonas*, a potentially opportunistic pathogen and spoilage organism, was a core member of the microbial community in the ileum [49,50]. The highest-scoring hub in the hindgut, *Acinetobacter,* is commonly present on beef, and there have been several reports of antimicrobial resistance genes in *Acinetobacter* isolates from beef samples [51,52,53]. When considering the intersection of pro-inflammatory, mucus-degrading bacteria and pathogenic proliferation, the host may be more prone to stress and systemic infection [46]. For instance, a higher prevalence of acid-producing bacteria can induce an epithelial inflammatory response and increase gut permeability [46]. Increased gut permeability is advantageous for certain pathogens that can migrate from the GIT into the blood stream [54]. For instance, *F. necrophorum* is highly associated with ruminal acidosis due to the increased opportunity to migrate from the rumen to the liver and form abscesses [54]. Therefore, understanding the impact of the microbial communities throughout the GIT is critical to modulate the negative effects of pre-harvest factors on food safety and meat quality.

The prevalence of dairy-beef cross cattle, the breed represented in this study, has been rapidly increasing, as reported by a 200% increase in the five-year average of beef semen sales in 2020, and presents significant economic benefits to dairy producers [55]. Despite this, dairy-beef crosses have a high prevalence of liver abscesses [56]. Liver abscesses have been associated with a decrease in average daily gain and hot carcass weight [57]. Therefore, while dairy-beef crosses offer significant economic benefits, they simultaneously present food safety concerns. In the present study, *Bacteriodes* was a core member in the duodenum, cecum, and large intestines, as well as the fifth most abundant genus in the cecum. *Bacteriodes,* has previously been identified in the liver abscess microbiome with *F. necrophorum*, supporting the hypothesized link between the GIT microbiome and liver abscess prevalence [57,58]. *Acinetobacter* has also been identified in the liver abscess microbiome, specifically in cattle fed tylosin phosphate [59,60]. Currently, tylosin phosphate, Tylan, is a popular feed additive in finishing diets to reduce liver abscesses. The cattle in this study were fed the Gain Master 55:35 RT #1707 pellet at a 5% inclusion rate, which contains Tylan (160g/ton) to prevent liver abscesses. Pinnell et al. (2023) [58] observed increased abundance of *Succinivibrionacae* UCG-001 in the rumen and *Turicibacter* in the colon epithelium when feeding tylosin, similar to digesta microbiome findings of this study. Despite reducing liver abscesses, tylosin has been observed to select for macrolide-resistant bacteria, posing a major food safety risk as common foodborne pathogens, such as *Enterococcus* species, *E. coli, Campylobacter*, and *Salmonella*, develop genetic resistance [61,62]. With growing concerns relating to liver abscess prevalence and antimicrobial resistance, modulating the microbiome has become a more desirable approach to controlling microbial pathogens in the GI tract of ruminants [61,62,58]. Therefore, future studies will be necessary to link the GIT microbiome at harvest with liver abscesses and antibiotic usage to determine future mitigation methods that reduce economic losses and improve food safety, especially in dairy-beef crossbred cattle.

The results of this study demonstrate future research strategies in harnessing the GIT microbiome to limit the welfare, meat quality, and food safety consequences of the pre-harvest process. For instance, protecting or modifying the rumen microbiome prior to harvest to improve resiliency may reduce the risk of liver abscesses or inflammation during feed withdrawal [54,63]. The small intestines epithelium has a high concentration of immune related cells, and by reducing the abundance of pro-inflammatory taxa under starvation conditions, such as *Streptococcus* or *Pseudomonas,* animals may experience less systemic inflammation [64]. Several studies have previously identified certain taxa in the feces, such as *Moryella* and *Clostridium,* to be associated with *E. coli* O157:H7 shedding in cattle [65,3]. Finally, while feed withdrawal has several positive benefits during the harvest process, its potential negative effects on the GIT microbiome, inflammation, and food safety may need further consideration. Additional research is necessary to better evaluate the pros and cons of feed withdrawal to balance ease of processing with food safety. The results of this study offer insight into the state of the entire cattle GIT microbiome at harvest.

## Acknowledgements

This research received no specific grant from any funding agency in the public, commercial, or not-for-profit sectors.

## Importance

With the global rise in antimicrobial resistance and the threat of foodborne illness, determining intervention strategies prior to harvest is a promising solution. The period between transportation from the feedlot to harvest may increase the risk of foodborne illness. During this period, cattle are withheld feed to reduce gastrointestinal tract (GIT) contents during carcass dressing. Feed withdrawal has many unintended consequences, such as acidosis and an increase in GIT pathogenic bacteria, that may result in foodborne pathogens on the final product. These consequences have yet to be thoroughly investigated in dairy-beef cross cattle, which have been rising in prominence in the United States. The GIT microbiome of dairy-beef cross cattle has been scarcely characterized, despite its influence on preventing the proliferation of common pathogens in the GIT. Therefore, it is necessary to determine the impacts of feed withdrawal on the GIT microbiome and its relation to foodborne illness.

